# Recombinant DNA resources for the comparative genomics of *Ancylostoma ceylanicum*

**DOI:** 10.1101/092494

**Authors:** Wadim J. Kapulkin, Adriana Magalska, Ewa Janecka, Arkadiusz Ciesielski, Malgorzata Lobocka, Jerzy M. Behnke, Halina Wedrychowicz

## Abstract

We describe the construction and initial characterization of genomic resources (a set of recombinant DNA libraries, representing in total over 90,000 independent plasmid clones), originating from the genome of a hamster adapted hookworm, *Ancylostoma ceylanicum*. First, with the improved methodology, we generated sets of SL1 (5‘-linker - GGTTAATTACCCAAGTTTGAG), and captured cDNAs from two different hookworm developmental stages: pre-infective L3 and parasitic adults. Second, we constructed a small insert (2-10kb) genomic library. Third, we generated a Bacterial Artificial Chromosome library (30-60kb). To evaluate the quality of our libraries we characterized sequence tags on randomly chosen clones and with first pass screening we generated almost a hundred novel hookworm sequence tags. The sequence tags detected two broad classes of genes: i. conserved nematode genes and ii. putative hookworm-specific proteins. Importantly, some of the identified genes encode proteins of general interest including potential targets for hookworm control. Additionally, we identified a syntenic region in the mitochondrial genome, where the gene order is shared between the free-living nematode *C. elegans* and *A. ceylanicum*. Our results validate the use of recombinant DNA resources for comparative genomics of nematodes, including the free-living genetic model organism *C. elegans* and closely related parasitic species. We discuss the potential and relevance of *Ancylostoma ceylanicum* data and resources generated by the recombinant DNA approach.

## Introduction

Hookworm infections are widely distributed in human and domesticated animal populations and pose a significant threat to global public health (Hotez 1995). The two anthropophilic species *Necator americanus* and *Ancylostoma duodenale* are responsible for most human infections, but hookworms of the genus *Ancylostoma* (i.e. *A. caninum, A. braziliense* and *A. tubaeforme)* also infect wild canids and felids, and are of importance in veterinary medicine through infections in small companion animals such as dogs and cats (Miller, 1971, 1979; Behnke 1990).

Unlike the two principal anthropophilic hookworm species, *Ancylostoma ceylanicum* predominantly infects wild as well as domesticated felids and canine hosts, but this species can also cause zoonotic infections in humans (Yoshida et al 1968; Areekul et al 1970; Behnke 1991;). The broad host range of *A. ceylanicum,* has allowed the establishment of a strain which can complete its life cycle in the golden hamster (Ray et al. 1972, Garside & Behnke 1989). This rodent adapted strain provides a unique opportunity for experimental studies on pathogenesis of hookworm infections in a convenient laboratory host (Menon et al. 1985, Behnke et al. 1997) and is a useful reference laboratory model for other hookworm infections and for studies that focus on host-immunity to hookworm infections (Behnke 1990, 1991, Alkazmi & Behnke, 2013). The major symptoms of hookworm infection, principally develop as a consequence of blood loss due to bleeding ulcerations and mucosal abrasions caused by adult worms browsing on the intestinal mucosa and feeding on blood from damaged blood vessels (Roche & Layrisse 1966, Kahn & Line 2010), resulting in an iron-deficiency anemia. Such lesions are also observed in the laboratory model of hookworm infection and are accompanied by other secondary signs i.e hypoproteinemia resulting from blood loss and weight loss (Garside & Behnke 1989). *Ancylostoma ceylanicum* infection induces immunological responses that have been shown to be protective in dogs and hamsters (Menon et al. 1985, Behnke et al. 1997), resulting in persistent acquired immunity and also in pups vaccinated with attenuated *A. caninum* larvae (Miller 1965).

Unlike the parasitic stages (activated L3 larvae, L4 and adults), the early larval stages that develop from eggs shed externally into the environment (L1, L2 and L3 larvae) are free-living. These early developmental stages are morphologically very similar in their microscopic appearance to the corresponding stages observed in *C. elegans.* Hookworms are evolutionarily and taxonomically closely related to the free-living nematode *C. elegans* (Blaxter et al. 1998), which is regarded widely as a highly informative and convenient manipulable laboratory model organism for a wide range of biological disciplines including developmental biology and genetics (Brenner 1974). *C. elegans* was the first metazoan animal to have its chromosomal genome sequenced (Sulston and Waterston, *C.elegans* Sequencing Consortium 1998) and its genome sequence is now considered a valuable reference in comparative analyses of gene and chromosome sequences of other nematodes, including hookworms. Moreover the transgenic technologies developed in *C. elegans* (reviewed in Mello & Fire 1995) have now been successfully exploited in developing recombinant genes from nematodes other than *C. elegans* (i.e. Kwa et al. 1995).

As part of our long term interests in comparative genome analyses of parasitic nematodes and also of our associated interests in the exploration of the potential of existing *C. elegans*– based transgenic technologies for the functional analysis of recombinant hookworm genes, we have constructed a set of the recombinant DNA libraries from the hamster adapted strain of *A. ceylanicum.* These recombinant DNA resources are described herein, as well as the methods that we used for the preparation of *E. coli* recombinants. We discuss the potential and relevance of data derived from work on *Ancylostoma ceylanicum* and the resources generated by the recombinant DNA approach.

## Materials and methods

### Isolation of nucleic acids and the recombinant library construction

Parasitic material was obtained from a strain of *A. ceylanicum* (referred as ‘Warsaw-strain’), maintained in Syrian hamsters for at least a hundred generations (the above hamster adapted strain was provide by Jerzy Behnke, University of Nottingham, UK, who originally obtained the strain in 1983 from Dr Rajasekariah of Hindustan CIBA-Geigy Ltd., Bombay, India. Hookworms were maintained in hamsters as described (Behnke et al. 1997). Adult worms were obtained from dissected intestines and harvested during necropsy. Pre-infective larvae were grown from synchronized eggs collected from dissected gravid females. Hatched larvae were kept in liquid media containing HBSS (Hanks’ balanced salt solution) at 25°C for two weeks. Nematode tissue samples were frozen at −80°C until further processing. The animal studies were approved by the Animal Care and Use Committee (R-508/98).

### Preparation and isolation of Nucleic acids

RNA isolation and SL1-cDNA preparation. mRNA was isolated using the Oligotex direct mRNA kit (Qiagen), from 20 adults or 2000 infective L3 larvae. Isolated poly A+ mRNA was converted to cDNA with *Thermus thermophilus* (rTth) DNA Pol I (reverse transcriptase activity), and PCR amplified using One-Step RT-PCR system (Gibco-BRL). PCR was carried out using the spliced leader (SL1 primer 5’-GGTTAATTACCCAAGTTTGAG) and oligo T (dT17) linker primers and the following amplification reaction profile: 45°C for 30 min, followed by 20 cycles of 94°C for 30 s; 55°C for 30 s; 70°C for 10 min followed by final extension 72°C for 30 min.

DNA isolation and preparation. DNA isolation was performed in agarose plugs according to the protocols of Osoegawa et al. (1998). Frozen adult worms tissues were shredded with micro dissecting scissors, inside a microfuge tube kept on ice. The contents of the tube were mixed 1:1 with melted 1% agarose in a 45°C water bath, then immediately poured into the acrylic glass (methyl methacrylate resin) McMaster chamber (0.15 ml). Solidified plugs were then extruded into the extraction buffer (2% SLS, 0.5M EDTA) supplemented with 2mg/ml of proteinase K (Sigma). Deproteinisation was continued at 50°C for 48h. After equilibration DNA was stored in 0.5M EDTA (pH 8.0) at 4°C. The quality of DNA was checked by analytical PFGE. Agarose embedded DNA was either electro-eluted into the dialysis bag as described (Strong et al. 1997), and sheared with a nebulizer and concentrated with an Amicon-50 spin dialysis filter (small insert genomic DNA library), or used directly for the construction of a Bacterial Artificial Chromosome library (according to protocols and vectors pBACe3.6 (Frengen et al. 1999) established by BACPAC-CHORI, Pieter de Jong personal communication).

Isolated nucleic acids (both RT-PCR amplified SL1-capped cDNA and sheared genomic DNA) were subsequently fractionated by preparative agarose gel electrophoresis. Briefly, after the preparative electrophoresis (45 min at 4°C at 5V/cm, TAE) the gels were sliced into three pieces and only two side fragments (containing molecular mass markers) were stained with ethidium bromide. Based on the marker bands the middle (unstained) part of the gel was sliced with subsequent vertical cuts and successive preparative fractions (a 0.3-0.5 cm of gel length) were taken covering the size range from 0.5-5.0kb). DNAs were released from the agarose gel matrix. Individual fractions were then used to build sub libraries. Both SL1-PCR cDNA and genomic DNA fractions were ligated in batches (containing fractions taken from the same preparative gel) into topoisomerase I based plasmid vectors [pCR-XL-TOPO™ and pCR-4blunt-TOPO™ vector (Invitrogen)] following the manufacturer’s instructions. 2μl of the ligation reaction was electroporated into freshly thawed aliquots of the electro-competent DH10B *E.coli* cells using the 1mm cuvette with a BTX ECM600 Electro Cell Manipulator. The resulting colonies were picked into 96 well plates containing LB media with 12.5% glycerol and stored at −80°C. Clones and libraries resulting from this work have been indexed,freeze-archived and distributed to the scientific community.

### Sequencing and sequence analysis

Sequencing reactions of library clones were outsourced to Microsynth Sequencing Service in Switzerland (www.microsynth.ch). In general in the first-pass sequence homology search, sequences scoring *E*-value of < 10^-5^ were considered as having significant homology. The sequences with homology to *C. elegans* proteins were identified by searching WormBase. Nucleotide sequence data without flanking primer sequences are available at dbEST/dbGSS GenBank™ under accession numbers (BI068132, BM536441-BM536500, BH739287-BH739307, BH739419) and analyzed with NCBI sequence suite (www.ncbi.nlm.nih.gov/genbank/). Where indicated individual sequences were conceptually translated into frames, and aligned with MAFFT multiple sequence alignment (mafft.cbrc.jp) with GUIDANCE confidence score (guidance.tau.ac.il). ER-routing signal sequence were identified on frame translated single letter coded string with SignalP (www.cbs.dtu.dk). GeneBank graphics or Ensembl BLAT/BLAST output representation were respectively used to generate graphical display of the relevant genomic annotations.

The sequences of the recombinant plasmids, were analyzed and searched against genome scale data sets. In the first step using the EST data from Mitreva et al. (2005) on *A. ceylanicum* SL1-ESTs (datasets generated using the SL1-cDNA libraries, we provided as material for sequencing) were aligned, to infer on allelic variants and correct for the first-pass sequencing and PCR errors. In the second step data from an *A. ceylanicum* scaffold assembly (Schwarz et al. 2015) <www.ncbi.nlm.nih.gov/biosample/SAMN02585415> were used to anchor the first pass sequence-tags within genomic scaffolds and infer on locally duplicated loci. The results of both steps of analysis with the genome scale data are summarized in Table 2.

## Results

### Recombinant DNA library construction

The four recombinant DNA libraries prepared from the hamster adapted *Ancylostoma ceylanicum* strain are summarized in the Table 1. The first two libraries are SL1 (spliced leader 1) PCR captured cDNA libraries and the remaining two libraries are genomic DNA libraries.

**Table 1.** The recombinant DNA libraries prepared from the hamster adapted *Ancylostoma ceylanicum.* pBACe3.6 is described in (Frengen et al. 1999).

| Library type | Nematode material | Cloning enzyme | Cloning vector | Insert range (kb) | *E.coli* colonies archived | Over - represented insert |
| --- | --- | --- | --- | --- | --- | --- |
| SL1-cDNA (PCR) | Infective L3 larvae | Vaccinia virus DNA topoisomerase | pCR-XL-TOPO™ | 0.25-10 | 20 000 | *mlc-5* (T12D8.6) homologue |
| SL1-cDNA (PCR) | Adults | Vaccinia virus DNA topoisomerase | pCR-XL-TOPO™ | 0.25-3.5 | 20 000 | Cysteine proteinase (F57F5.1) homologue |
| Genomic DNA | Adults | Vaccinia virus DNA topoisomerase | pCR4blunt-TOPO™ | 2-10 | 20 000 | Mitochondrial DNA |
| Genomic DNA | Adults | EcoRI / T4 DNA ligase | pBACe3.6 | 30-60 | 30 000 | - |

The cDNA libraries represent *E. coli* colonies electro-transformed with ligated PCR products. The PCR products were derived from synthesis of the poly-adenylated mRNA material isolated from infective L3 larvae and adult nematodes (respectively), using the the Tth DNA polymerase and oligo-dT and 5’ SL1 primers. The PCR products robustly amplified the material, typically rendering the fairly uniform range of products (0.25-10kb and 0.25-3.5kb, respectively for larval and adult isolated material) as visualized by agarose gel electrophoresis (not shown). Provided the SL1-dT PCR products rendered intensively amplifying bands (apparent after the gel electrophoresis), we sized the gel fractionated material by horizontal slicing of the gel after the preparative electrophoresis. This procedure was implemented as an improvement in the quality of the constructed libraries and intended to remove the over-amplifying by-products from the PCR material. The material extracted from consecutive gel-sized fractions was subsequently ligated in 6 and 4 (respectively for larval and adult extracted material, were fraction third and second were respectively labeled as containing the over-amplifying bands) independent reactions and later treaded as separate library segments.

The genomic DNA libraries were prepared according to established protocols, using the agarose-gel embedded/extracted DNA and gel fractionation following DNA electroelution (Osoegawa et al. 1998). The blunt-end small insert library was prepared from mechanically sheared DNA molecules while the BAC library was prepared using the DNA digested with EcoRI restriction enzyme.

### Recombinant DNA library characterization

The initial characterization of the prepared libraries was performed by the EcoRI restriction enzyme digestion of isolated plasmid DNAs. To evaluate plasmid insert integrity and diversity the EcoRI digestion products were size compared using the estimates from routine agarose gel analysis (not shown). In this step, we immediately confirmed the cDNA libraries were >95% recombinant and therefore concluded that the prepared collection was of adequate quality. This conclusion was based on: first on size polymorphism of individual plasmids, and secondly on presence of internal EcoRI sites. At this stage we confirmed the two amplified SL1-cDNA fractions, corresponding to fraction three and two (respectively for larval and adult extracted material) originally labeled as containing the over-amplified PCR products lacking the required EcoRI size polymorphisms (not shown). In the case of plasmids prepared from colonies grown for the blunt-end small insert genomic library we could not confirm the same level of integrity. Instead we observed that the overall rate of ~75% recombinants was retrieved across all fractions tested. In contrast plasmids grown for the colonies of the transformed BAC library were of the expected size at 30-60kb and were ~95% recombinant.

In order to confirm the estimates from the agarose gel analysis we subjected the initial preparations to automated Sanger DNA sequencing (this was done by the outsourcing to the commercial DNA sequencing service). Analysis of the sequencing products is summarized in the Table 2. We initially proceeded with six plasmids per SL1-cDNA fraction. This step resulted in sequence based identification of the over-amplified PCR products (listed with Table 1. as over represented inserts.) avoiding the potentially problematic segments of libraries. In the case of the adult SL1-cDNA library the identified over-amplified insert corresponds to *A. ceylanicum* cysteine proteinase. The particular *A. ceylanicum* cysteine proteinase has been reported by our group previously (Mieszczanek et al. 2000) using RACE PCR. In the case of the larval SL1-cDNA library the over-amplified product corresponds to *A. ceylanicum* protein homologous to *mlc-5 C. elegans* gene. Those unwanted and non-polymorphic segments (where in each case the sequencing of six out of six attempted clones confirmed the same cDNA sequence) were subsequently excluded from the set of freeze archived colonies.

**Table 2.**
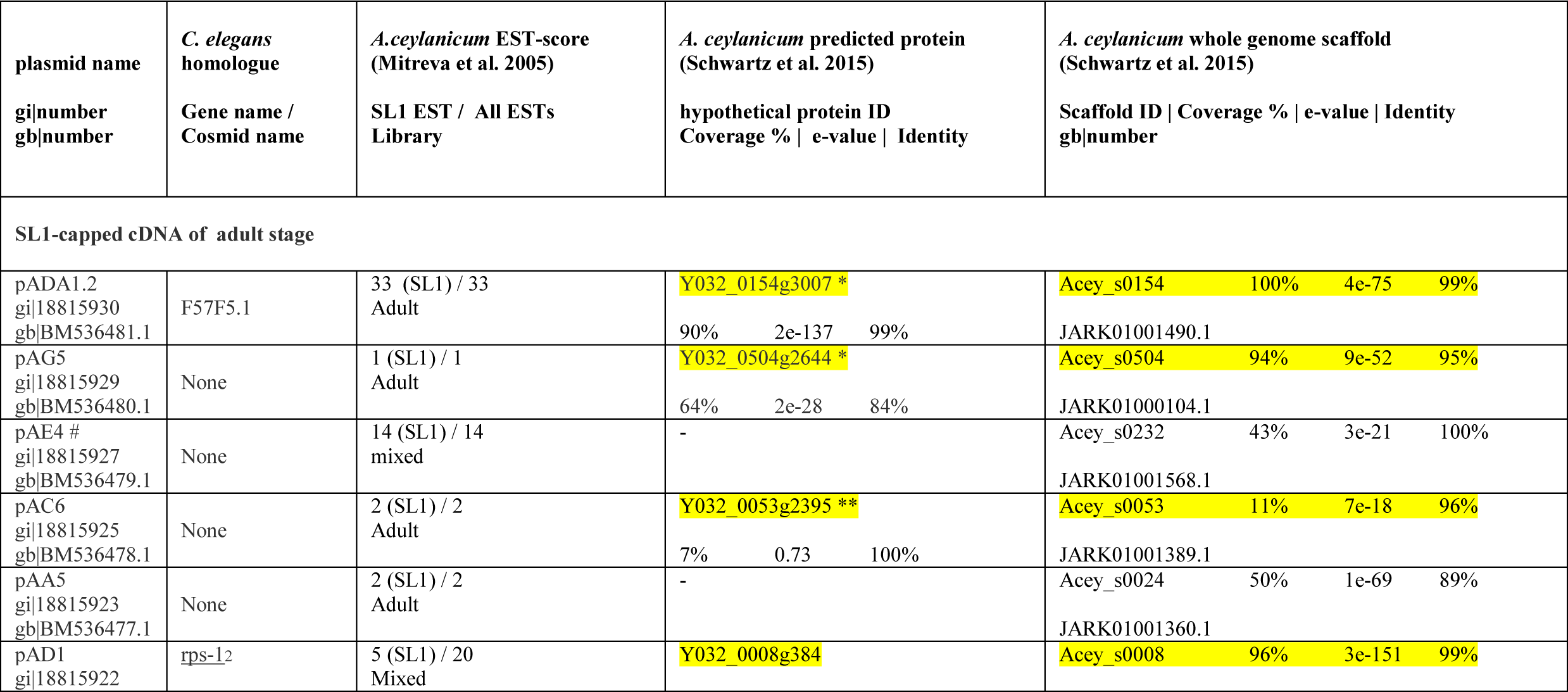
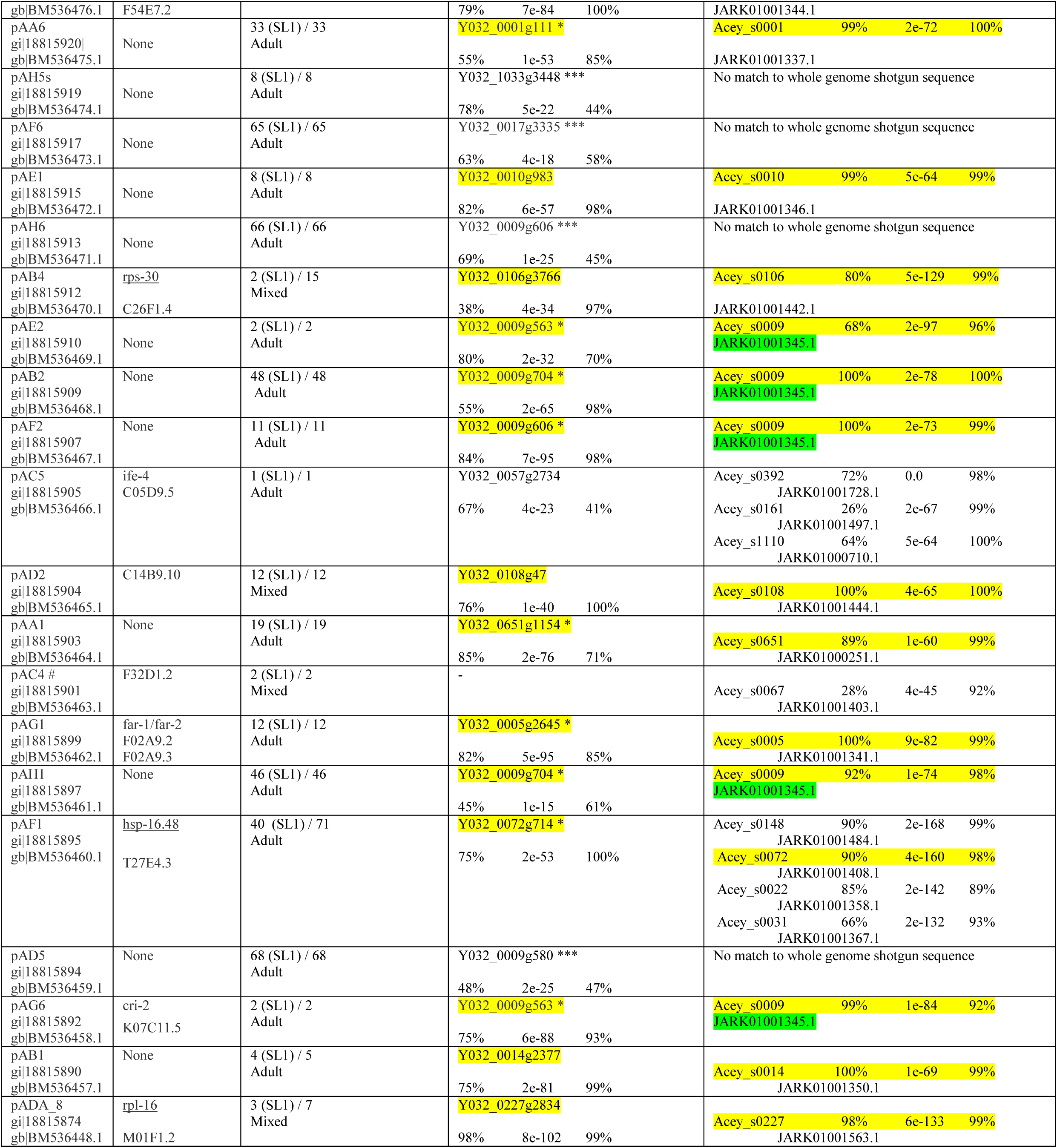
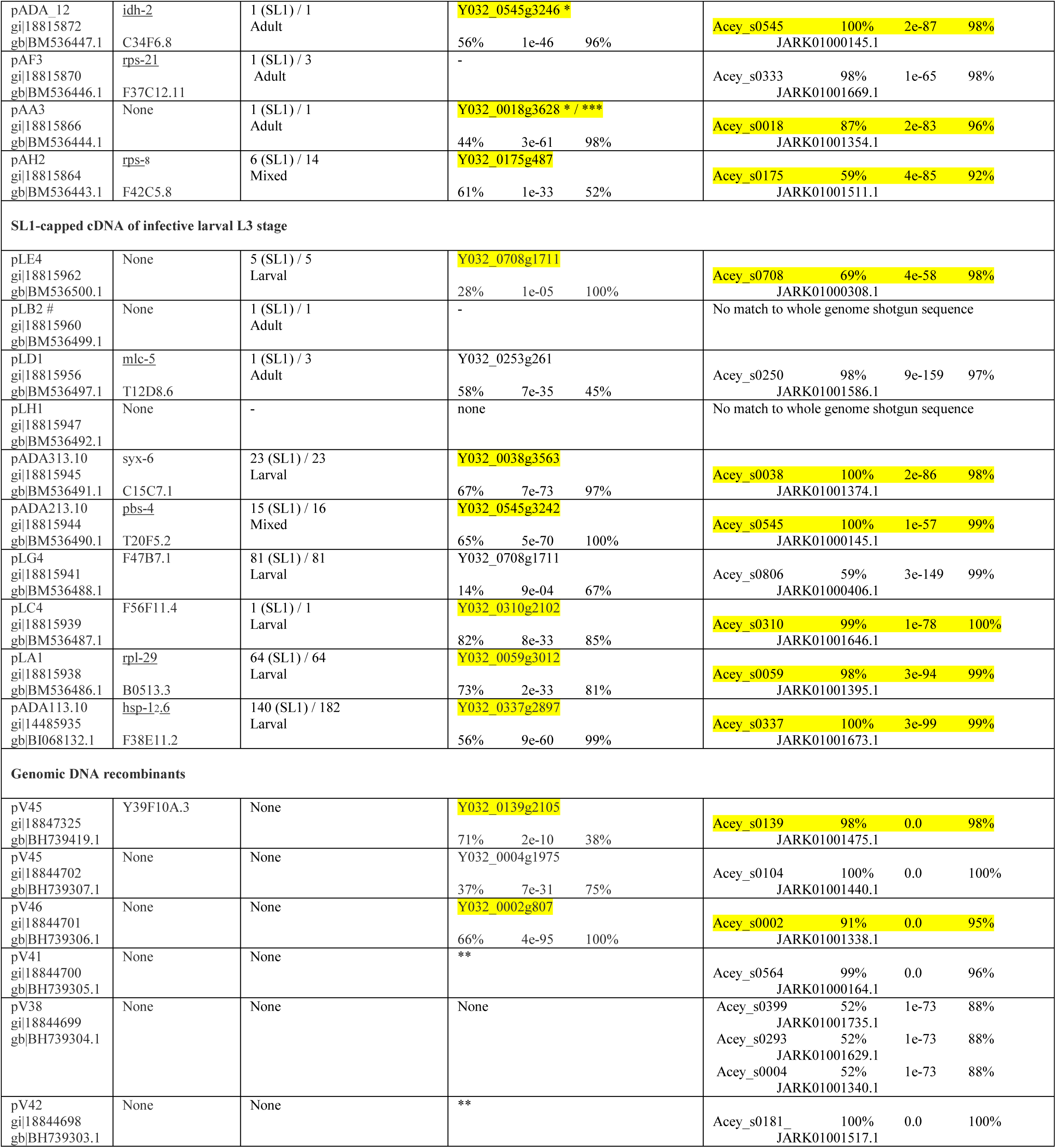
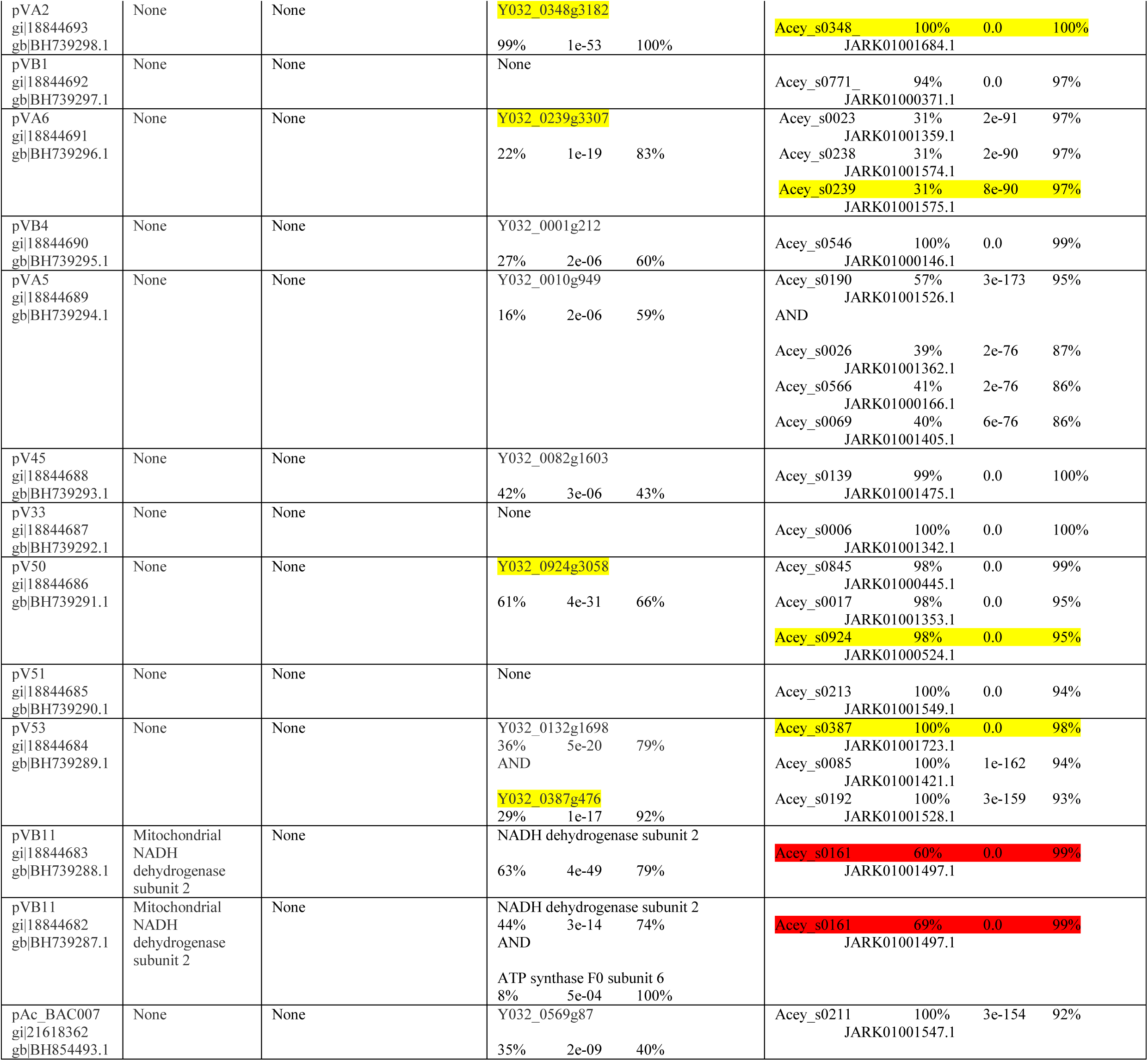
Column 1^st^: *E. coli* plasmid clone name and GeneBank (short reads are indicated by #).; Column 2^nd^: *C. elegans* homologue gene name and database identifier (cosmid name). Column 3^rd^ *A. ceylanicum* EST-score. Number of ESTs re-sampled from SL1-cDNA library / all EST reported by (Mitreva et al. 2005) and library-of-origin (infective L3 and / or adult). Column 4^th^: hypothetical proteins predicted by assembly described in (Schwartz et al. 2015) supported by conceptually translated homology based search. Best protein match is typically shown (except for pV53 where two non overlapping regions of homology were detected across single sequencing read and reads from pVB11 where two non overlapping regions of homology were detected across single sequencing read derived from BAV82618.1), with following disclaimers: ‘*’ indicates where other protein matches were predicted on the same scaffold i.e. in case pADA1.2 cysteine proteinase homologues, indicating locally duplicated gene clusters, possibly forming multi-allelic loci; ‘**’ match below significance threshold, concordant with scaffold ID; ‘***’ indicates where other protein matches were predicted on the different scaffold i.e. Acey 0009 scaffold;. Column 5^th^: unique matches are shown when possible, otherwise top three scaffold matches are listed (except for pAF1 hsp-16 homologues, where first 4 putative candidates are listed and pVA5, where two non overlapping regions of homology were detected across single sequencing read, specifying a single scaffold Acey_s0190 on the left and three other scaffolds on the right). Positive hypothetical protein identifications are highlighted in yellow in cases if they match with the identified scaffold (total 33 tentative pairs). Hypothetical proteins predicted on the Acey 0009 scaffold, are highlighted in green. Mitochondrial segment pVB11 sequences identified as matching the scaffold Acey 0161, with high confidence are highlighted in red.

The results of the sequencing of plasmids from the genomic DNA libraries are included in Table 2. In addition to a lower than expected fraction of recombinant plasmids from the blunt-end small insert genomic library (above) we observed that some of the *E. coli* colonies grew poorly (with apparent growth delay in over-night mini-cultures) resulting in lower quality templates for sequencing reactions. These colonies were verified as containing the rearranged vector sequences (~25%) and were excluded from further analysis. However, despite the above difficulties, we managed to generate informative reads of truly recombinant plasmids from this library. Notably, with sequence based characterization we identified a plasmid corresponding to a segment of *A. ceylanicum* mitochondrial DNA. This result indicated that the fraction of the blunt-end small insert library is contaminated with non-nuclear *A. ceylanicum* DNA. At this stage we attempted to generate paired-end sequences of the BAC library clones, however at the time the sequencing reactions were attempted, of six confirmed EcoRI polymorphic clones, sequencing reactions resulted in only a single informative read (conducted at the Institute for Biochemistry and Biophysics-PAS). Because of limited resources, as well as a restricted time frame for the project,we were forced to abandon the further characterization of the constructed and freeze archived *A. ceylanicum* BAC library.

### Sequence based identification of the evolutionary conserved nematode factors

Based on the *A. ceylanicum* sequences generated during the initial characterization of the recombinant DNA library, we sought confirmation of the presence of the evolutionarily conserved factors i.e. genes present in nematodes other than hookworms. We therefore first compared the generated sequences to genes known to exist in the model organism *C. elegans* (Table 2.). In addition to the two over-represented factors mentioned above (Table 1.) we identified the following *C. elegans* homologues: rps-12, F54E7.2; rps-30, C26F1.4; ife-4, C05D9.5; C14B9.10; F32D1.2; far-1/far-2, F02A9.2/F02A9.3; hsp-16.48, T27E4.3; cri-2, K07C11.5; rpl-16, M01F1.2; idh-2, C34F6.8; rps-21, F37C12.11 and rps-8, F42C5.8 in adult SL1-cDNA library and syx-6, C15C7.1; pbs-4, T20F5.2; F47B7.1; F56F11.4; rpl-29, B0513.3; hsp-12.6, F38E11.2 in infective L3 SL1-cDNA library and Y39F10A.3 in the *A. ceylanicum* genomic DNA library.

If any of these conserved factors were later reported by others in descriptions of material from our libraries (Mitreva et al. 2005), we do not here repeat analysis of relevant genes. However we have attempted to retrospectively identify which of the above conserved factors are most prevalent when sampled with the larger SL1-cDNA set (Mitreva et al. 2005). Based on the data included in Table 2, we have weighted the relative abundance of individual factors by counting absolute numbers of the SL1 ESTs generated by our collaborators. We have established that small heat shock proteins exceed in number other factors, with homologues of *C. elegans* hsp-12.6 (F38E11.2) protein and hsp-16.48, T27E4.3 accounting for 140 SL1 ESTs and 42 uncapped ESTs and 40 SL1 ESTs and 31 uncapped ESTs in larval and adult libraries respectively. Interestingly the identified homologue of F47B7.1, poorly characterized *C. elegans* gene of uncertain function appears also highly represented (accounting for 81 SL1 ESTs found exclusively amongst clones from the library we provided). For comparison abundant ribosomal protein rpl-29, B0513.3 is represented by 81 SL1 ESTs amongst sequences derived from clones from ours SL1-cDNA library. Table 2 lists additional homologues of *C. elegans* genes derived from clones from the SL1-capped cDNA libraries we provided, but strikingly absent from uncapped libraries provided by other collaborators. This class of SL1-capped *A. ceylanicum* genes includes homologues of the following *C. elegans* genes: C14B9.10 and pbs-4 (supported by respectively 12 and 16 combined SL1 ESTs identified in both adult and L3 libraries); far-1/far-2 and cri-2 (supported by respectively 12 and 2 SL1 ESTs identified specifically in adult SL1-cDNA library) and syx-6 (supported by 23 SL1 ESTs identified specifically in larval SL1-cDNA library). This analysis has further revealed that some conserved factors identified by our pilot analysis as homologous to predicted *C. elegans* genes (ife-4, C05D9.5; idh-2, C34F6.8 and F56F11.4) were apparently not sampled with the larger set (Mitreva et al. 2005). We note specifically that the current description is so far the only report on those conserved trans-spliced, nematode factors present in *A. ceylanicum.*

### Sequence based identification of the taxonomically restricted genes encoded by the *A. ceylanicum* genome

In contrast to the above evolutionarily conserved nematode factors, where sequence homology with known *C. elegans* genes (and often with other predicted nematode genes other than caenorhabdids, typically including genes described in hookworms other than *A. ceylanicum*) could be established, within the set of the initial sequences derived from the recombinant libraries, we identified a number of predictions where homology to known *C. elegans* genes could not be inferred. Provided the genome of *C. elegans* is complete, and all predicted sequences have been deposited in public databases, we regard the other sequences derived from our libraries (examples where homology to *C. elegans* genes could not be established) as taxonomically restricted. Those taxonomically restricted sequences contributed to approximately half (20 of 39 sequences derived from the combined larval and adult SL1 capped cDNA libraries) of the sequences we generated while characterizing the recombinant DNA libraries. The above trend is more prominent in the SL1 capped adult set (13 of 30 unique sequences) when compared to larval SL1 set (7 of 9 unique sequences) but unexpectedly absent from the control set of genomic sequences (where there is only one sequence matching the predicted *C. elegans* gene Y39F10A.3 amongst 19 unique genomic reads).

In view of the above set of the taxonomically restricted genes encoded by *A. ceylanicum,* in the larger sampling of our SL1-capped cDNA libraries, we again attempted to retrospectively identify which of the taxonomically restricted factors are most prevalent in the larger SL1-cDNA set (Mitreva et al. 2005). We therefore repeated the type of analysis we had conducted for hookworm genes evolutionarily conserved with *C. elegans* and weighted the relative abundance of individual taxonomically restricted genes by counting the absolute numbers of the SL1 ESTs generated by our collaborators (Table 2). Based on the above analysis we observed that again certain SL1-capped transcripts of *A. ceylanicum* (denoted here as plasmid clones pAG5, pAE2 and pAA3) were apparently not sampled in the larger set described by (Mitreva et al. 2005). The above three examples of adult SL1-capped cDNAs (pAG5, pAE2 and pAA3) were assigned as homologous to different sets of hypothetical proteins / protein families of *A. ceylanicum* (represented by Y032_0504g2644, Y032_0009g563 and Y032_0018g3628 respectively, based on the relative homology score, sequence coverage and percent identity in the aligned region), presumably expressed from at least three different genomic loci (Table 2). Strikingly, those homologues of the three hypothetical proteins predicted in *A. ceylanicum* appeared taxonomically restricted i.e. where the sequence homologies were strictly species specific or where the only homologues were identified in hookworms other than *A. ceylanicum.* Based on this analysis, we interpret these unique SL1-capped ESTs as representing adult worm enriched trans-spliced transcripts of relatively low abundance. We note specifically that this description is so far the only report on these taxonomically restricted trans-spliced factors recovered from adult stages of *A. ceylanicum* identified explicitly by our pilot analysis of clones derived from the recombinant DNA resources. The above analysis was further extended into the sequences derived from other plasmid clones prepared in parallel during the initial characterization of our *A. ceylanicum* recombinant DNA resources. This analysis confirmed further that the additional examples of SL1-capped transcripts identified in our preliminary characterization were specifically recovered from the libraries we provided for the more extensive sampling reported by our collaborators (Mitreva et al. 2005). Of the factors listed in Table 2, based on the assignment guided by the homology search, we further included the following plasmid clones (pAA6, pAF6, pAE1, pAH6, pAB2, pAF2, pAA1, pAH1, pAD5 and pAH5 homologous to *A. ceylanicum* hypothetical protein / protein families represented by Y032_0001g111, Y032_0017g3335, Y032_0010g983, Y032_0009g606, Y032_0009g704, Y032_0009g606, Y032_0651g1154, Y032_0009g704, Y032_0009g580 and Y032_1033g3448 respectively) into the above group of taxonomically restricted factors derived exclusively from SL1-capped cDNAs recovered from adult stage of *A. ceylanicum.* We note that these ten SL1-capped transcripts represent novel proteins of mostly unknown function presumably specific to the adult stage of *A. ceylanicum.* However in contrast to the SL1-capped transcripts uniquely defined by our pilot characterization, the above group of ten factors is supported by the more extensive sampling reported by Mitreva et al. (2005). Specifically, the dataset described by our collaborators includes further examples of SL1-capped transcripts corresponding to the above adult specific factors (supported by 33, 65, 8, 66, 48, 11, 19, 46, 68 and 8 trans-spliced cDNAs respectively) suggesting a pattern typical for abundantly expressed transcripts. In the particular examples of SL1-capped cDNAs from plasmid clones pAF6, pAH6 and pAD5, these were represented by >60 ESTs in the larger set of sampling from the library we provided, suggesting that those exceed in relative numbers other transcripts recovered from adult stage. Our analysis suggests that the proteins defined by the above abundant transcripts are likely hookworm or even species specific and that they perform presumably relevant (and collectively essential) functions in the parasitic adult stages *A. ceylanicum.* We suggest therefore that the above sampling of the SL1-capped trans spliced cDNAs are biased toward taxonomically restricted factors expressed in the adult stage of *A. ceylanicum.* This perspective appears plausible in the context of the two other examples of SL1-capped transcripts identified in the preliminary characterization of recombinant DNA clones from our libraries. First, sampling from the SL1-capped adult library included a sequence derived from cDNA clone pAB1 (supported by 4 SL1-capped ESTs derived from the library that we provided and one EST derived from uncapped cDNA library provided by other collaborators). pAB1 encodes for factor markedly homologous to *A. ceylanicum* hypothetical protein Y032_0014g2377, which in contrast to the above factors appear taxonomically unrestricted. While factor encoded by pAB1 appears taxonomically excluded from caenorabdids (paraphyletic with respect to), predictions homologous to *A. ceylanicum* hypothetical protein Y032_0014g2377 could be found in *Necator americanus* (NECAME_00160), *Dictyocaulus viviparous* (DICVIV_13749), *Oesophagostomum dentatum* (OESDEN_11368) and *Haemonchus contortus* (CDJ80097.1, unnamed protein product)(not shown). Second, in our sampling there appears to be the only larval match (supported by two trans spliced cDNAs pLE4 and pLC2, independently recovered from the infective L3 larvae SL1-capped library we provided and represented further by total 5 SL1-capped ESTs in larger set of samplings reported by ours collaborators in 2005) representing a potential example of a taxonomically restricted gene present in a non-adult stage. We note however in the latter case that confidence appears low and that this is due to a very limited (below the threshold of the relative homology score) homology to the predicted *A. ceylanicum* hypothetical protein Y032_0708g1711.

### The clues retrieved from mapping of the *A. ceylanicum* recombinant DNA clones onto the scaffolds corresponding to the genome sequence

While conducting the previous steps of the analysis on the sequences derived from the preliminary characterization of the recombinant DNA resources, we have observed that sequences derived from certain clones could not be discretely assigned into different clusters in an unambiguous manner using the available cDNA samplings from Mitreva et al. (2005). We have observed that a subset of sequences derived from apparently different trans-spliced transcripts from the adult SL1-capped cDNA library, appear mutually homologous in sequence based searches. This behavior is indicative for transcripts encoded by multi-allelic loci or transcripts encoded by the gene families (i.e. *C. elegans* small heat shock proteins) and becaue of the absence of a draft of the genomic sequence, this poses a particular challenge for taxonomically restricted genes and reads from clones derived from genomic libraries. Among the highly expressed taxonomically restricted genes that we initially noted, three different SL1-capped cDNA clones pAB2, pAH6 and pAF2 were mutually homologous. Using those sequences as specific identifiers, with a re-iterated search, we established that three additional sequences derived from clones pAD5, pAF6 and pAH1. generated during our preliminary characterization attempt. could be assigned into the same homology clade. Subsequently using SL1-capped cDNA sequences deposited in public databases, we identified further pAA3, pAH5 and pAA6 by aligning with overlapping sets of *A. ceylanicum* predicted proteins. We termed those groups of predicted *A. ceylanicum* proteins LKIN-motif family proteins, after the distinct Leu-Lys-Ile-Asn tetrapetide motif which occurred in the predicted sequence of the first discovered member derived from SL1-cDNA pAB2 (Fig 1. and 2.). In order to assign the individual members of the LKIN-family into individual loci we took advantage of the scaffold-based assembly corresponding to the *A. ceylanicum* genome sequence deposited in a public database by Schwartz et al. (2015). The results of mapping of the sequences generated during the preliminary characterization of the recombinant DNA resources onto *A. ceylanicum* genomic scaffolds described in Schwartz et al. (2015) are summarized in the last column of the Table 2.

**Figure 1.**
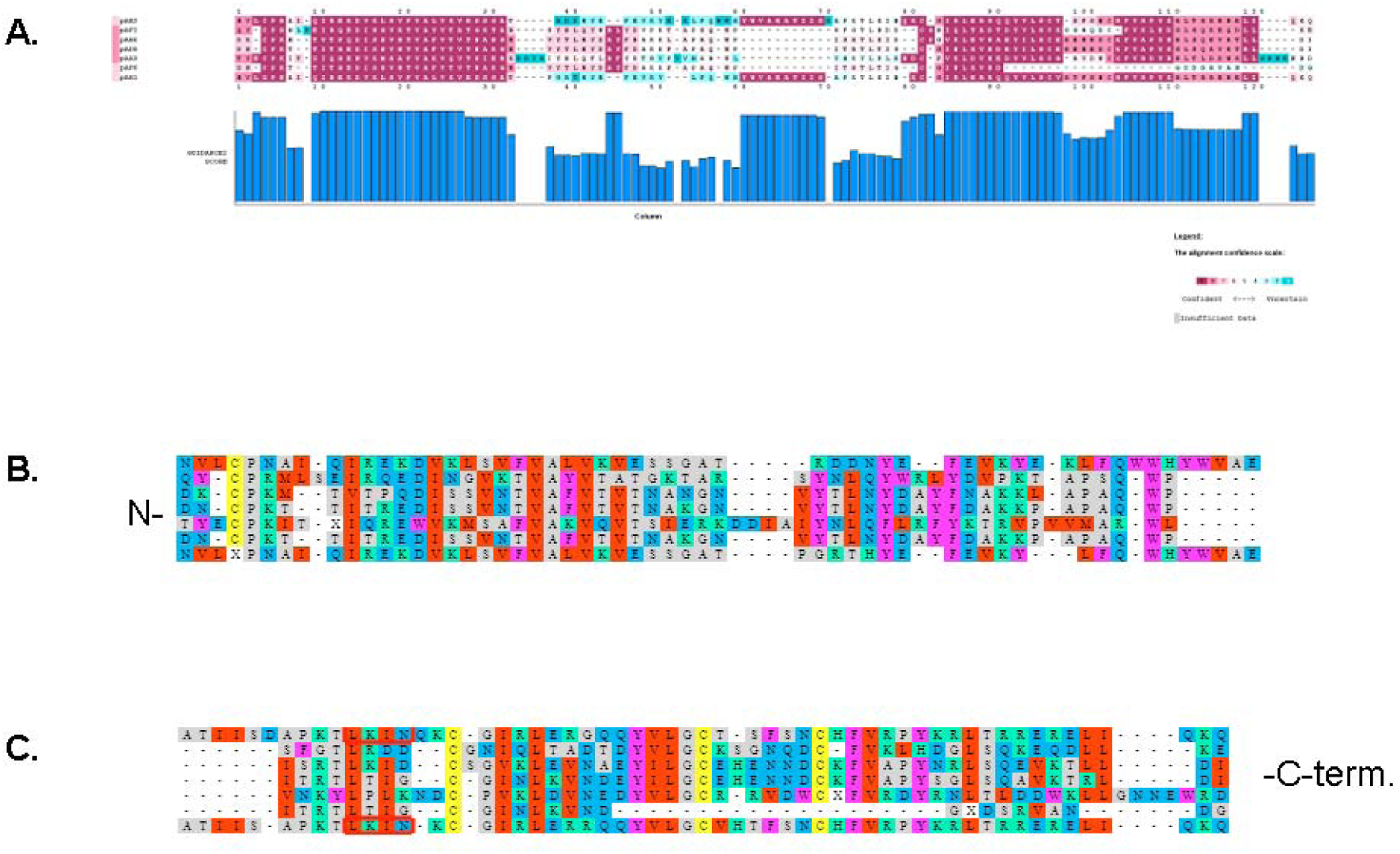
Color coded alignment of LKIN-motif proteins identified based on recombinant SL1-capped cDNA clones described in the study. Sequences deduced from the first seven SL1-capped clones of adult *A. ceylanicum* library are shown. Sequences are given in the following order: pAB2, pAF6, pAH6, pAD5, pAA3, pAF6 and pAH1. The top four (pAB2, pAF6, pAH6, pAD5) representing conceptually translated sequences are included as directly deduced from high quality reads. The remaining three (pAA3, pAF6, pAH1) include uncertain bases (masked with X). pAF6 may also include an artifact and represent truncation of the actual sequence. The pAH1 sequence derived from low quality reads were edited to remove frame-shifts. All sequences are presented as mature peptides (SignalP predicted N-terminal ER routing signal is trimmed). A. MAFT-7 multiple sequence alignment color coded by GUIDANCE confidence score of LKIN-motif proteins < guidance.tau.ac.il>. B. The N-terminal part of the detailed colored alignment, C. The C-terminal part of the detailed colored alignment. Note that the LKIN-motif in pAB2 and pAH1 is framed in the red box.

**Figure 2.**
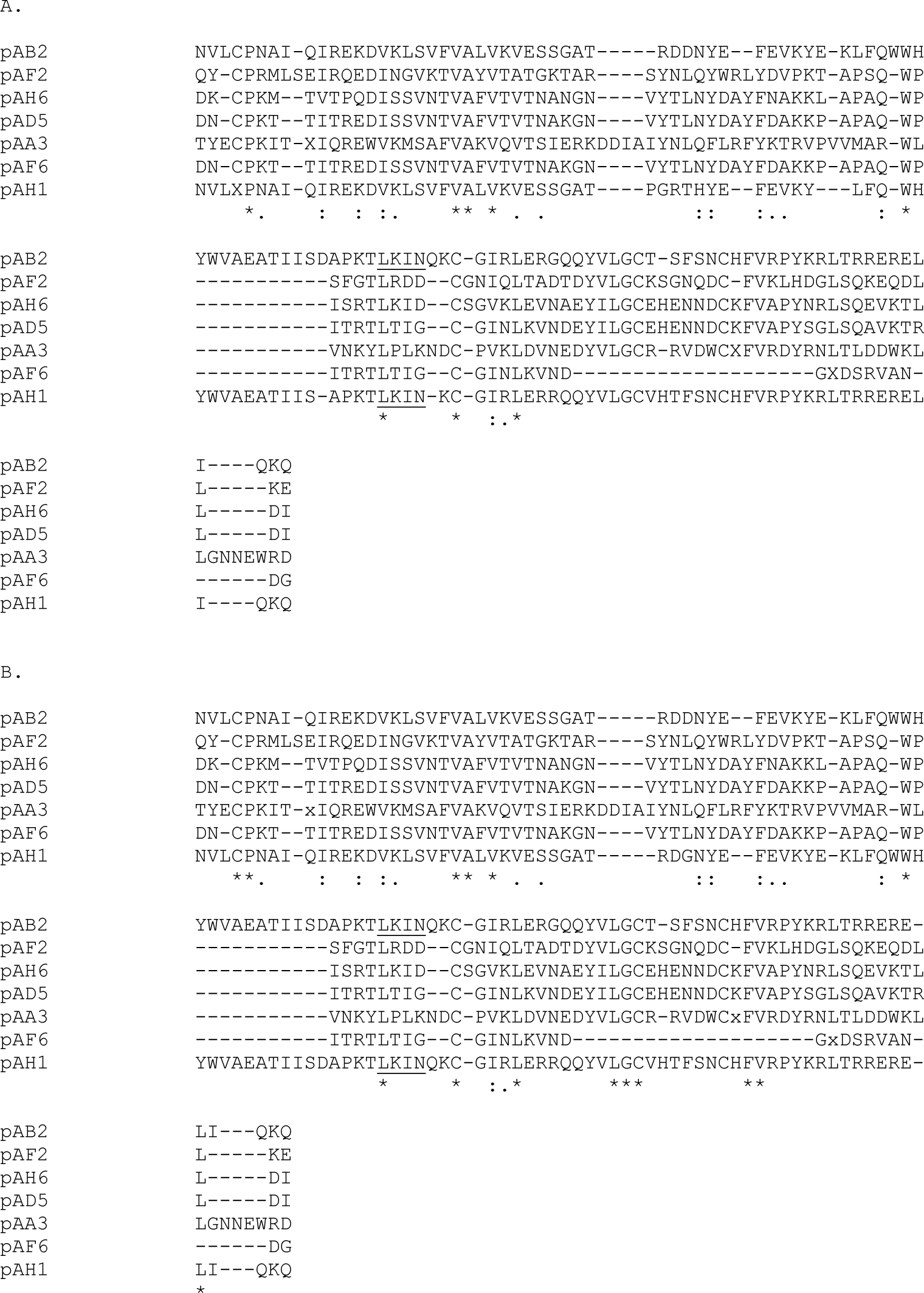
LKIN-motif proteins identified in the study, presented as CLUSTAL format alignment by MAFFT (v7.306b) of sequences presented with Fig.1., generated by ‘G-INS-1’ method (Katoh et al. 2005) < mafft.cbrc.jp/alignment/software>. A. Original MAFFT-7 alignment. B. MAFFT-7 alignment, manually edited, based on SL1-capped EST (`pj15c02.y1`) concordant with pAH1. Residues conceptually translated from ambiguous bases of first pass reads are coded as lowercase ‘x’. LKIN-motif is underlined.

Based on the above scaffold-mapping data analysis that we generated, we then retrieved the genomic coordinates aligning to the sequences we generated during the preliminary characterization stage, and initially focused on SL1-capped cDNAs encoding for the LKIN-motif family members. We have established that two of the LKIN-motif family members (represented here by SL1-capped cDNA clones pAB2 and pAF2) specify duplicated pairs of loci linked on single genomic scaffold Acey_s0009 (JARK01001345.1) (Figure 3.). This analysis has identified SL1-capped cDNA clone pAB2 as encoding two predicted proteins Y032_0009g704 and Y032_0009g716 as encoded by the locally duplicated pair of genes, predicted at positions Acey_s0009.g704-JARK01001345.1[2,052,912..2,053,464] and Acey_s0009.g716-JARK01001345.1[2,086,709..2,097,446]. We termed that pair of genes the LKIN-motif family locus AB2. Next, we have established the second identified SL1-capped cDNA clone pAF2 specified by two predicted proteins Y032_0009g606 and Y032_0009g591 as encoded by the second pair of presumably duplicated genes, predicted at positions Acey_s0009.g606-JARK01001345.1[1,423,074..1,425,309] and Acey_s0009.g591-JARK01001345.1[1,399,607..1,400,045]. We refer to that pair of genes as the locus AF2. Based on genomic coordinates, the genomic position of two locally duplicated gene pairs spanning a < 0,7 MB interval, located in the middle of the single genomic scaffold Acey_s0009, constitutes the locus encoding for the major class of the abundant SL1-capped transcripts specific to adult stage of the *A. ceylanicum.* Conducting the same type of analysis, with the other clones corresponding to the above class of adult specific SL1-capped transcripts, revealed that unlike clone pAH1 (supporting the LKIN-motif locus AB2), clones pAD5, pAH6, pAF6 and pAH5 could not be assigned to specific scaffolds described by Schwartz et al. (2015) despite apparently encoding for proteins homologous with other members of the LKIN-motif family of proteins (Table 2.). However, the homology based search, indicated that those adult specific SL1-capped cDNA were predicted to support the placement of the LKIN-motif family locus on the single genomic scaffold Acey_s0009. Moreover, the same search indicated that around and within the specified 0,7 MB interval there are apparent (albeit more distant) additional genes encoding for hypothetical proteins homologous with the LKIN-motif family members. Specifically, those gene predictions supported the additional hypothetical proteins encoded by the predicted genes grouped in the proximal (referring to scaffold coordinates) part of Acey_s0009. Y032_0009g537, Y032_0009g542, Y032_0009g544, Y032_0009g545, Y032_0009g580, Y032_0009g585 and Y032_0009g586 were positioned outside of the locus AF2 (Fig. 3. A and B), while Y032_0009g594 was predicted to lie between Y032_0009g591 and Y032_0009g606, and is therefore positioned internally within the AF2 locus (Fig. 3. C). Next, genes encoding for hypothetical proteins Y032_0009g610, Y032_0009g612, Y032_0009g678, Y032_0009g683, Y032_0009g686 are predicted to span the interval between AF2 and AB2 loci (Fig. 3. D and E) and Y032_0009g705 was predicted to lie between Y032_0009g704 and Y032_0009g716 and therefore positioned internally within the AB2 locus (Fig. 3. F and G). Collectively, by conducting repeated searches we have confirmed at least 18 distinct predicted genes as encoding for adult specific, diverse protein LKIN-motif members, all being predicted to occur within the defined interval (~1MB in length) of the single genomic scaffold Acey_s0009. Provided this type of distribution is characteristic for gene families expanded by local duplications, we have established that the genomic scaffold Acey_s0009 encodes for a hookworm specific multi-allelic locus. In this phase of the analysis we concluded that some of the most abundant hookworm specific SL1-capped transcripts, encoding for highly diverged and presumably secreted proteins (based on the N-terminal SignalP predictions of the ER-routing signal, not shown) are likely to be encoded by clustered genes that have arisen through local gene duplications.

**Figure 3.**
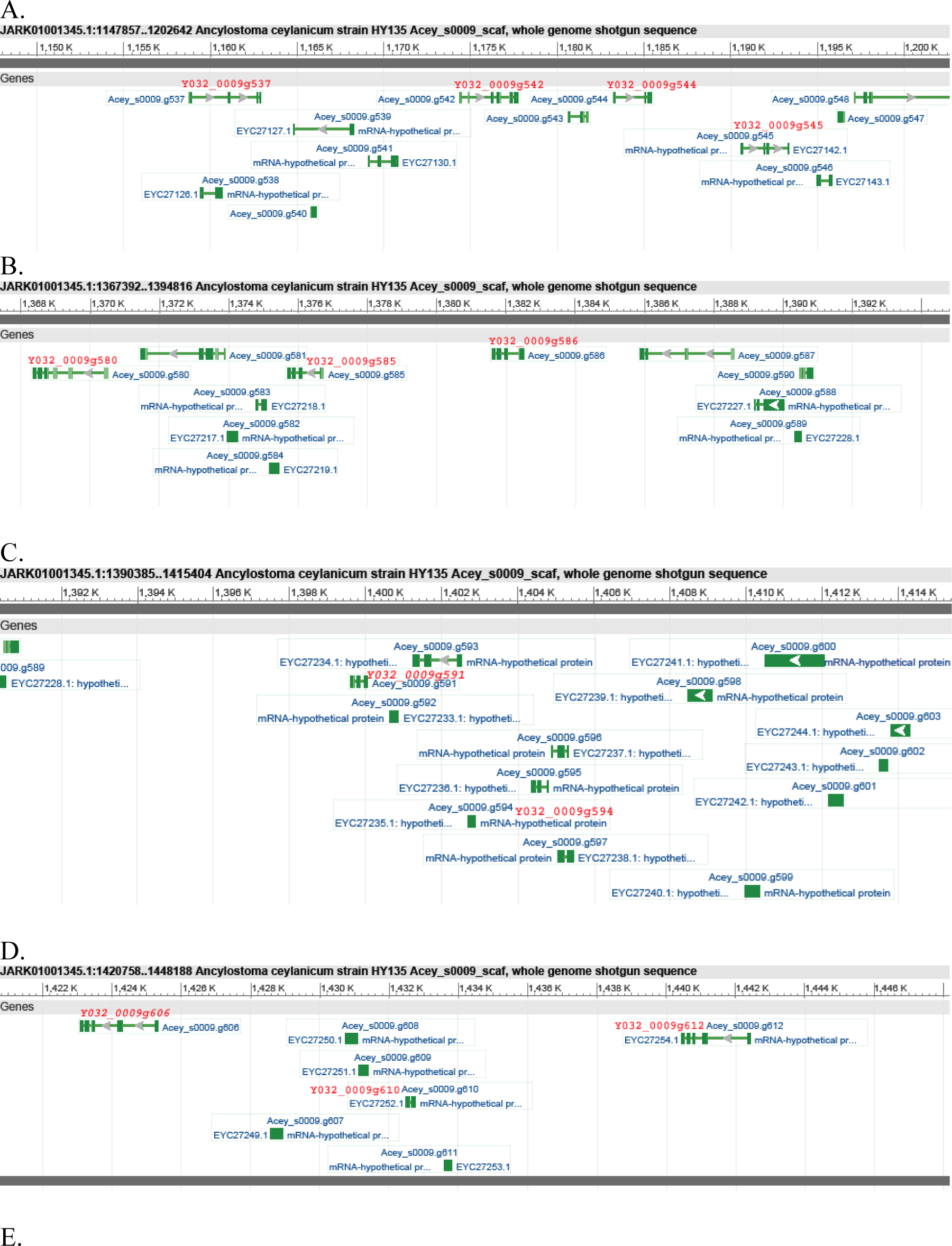
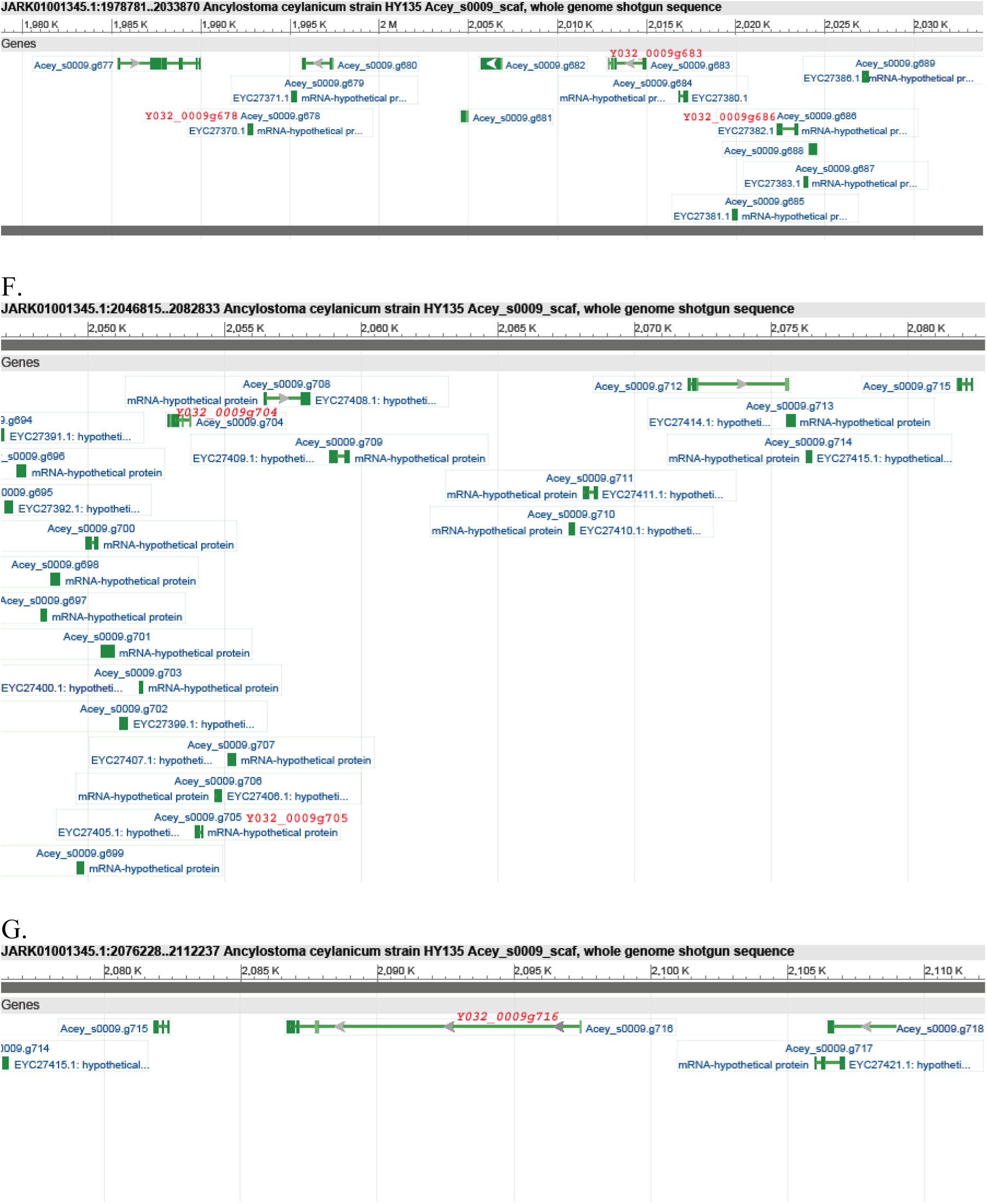
LKIN-motif protein family locus architecture on Acey_s0009 scaffold. LKIN-motif family protein names are written in red font, above the predicted genes represented by GeneBank graphics. Proteins predictions directly supported by the SL1-capped cDNA pAB2 (Y032_0009g704 and Y032_0009g716) and pAF2 (Y032_0009g591 and Y032_0009g606) are marked by red oblique font. A. LKIN-motif family proteins Y032_0009g537, Y032_0009g542, Y032_0009g544 and Y032_0009g545 identified in the most proximal part of scaffold Acey_s0009 in region corresponding to JARK01001345.1 [1,147,858..1,202,642]. B. LKIN-motif family proteins Y032_0009g580, Y032_0009g585 and Y032_0009g586 identified in the proximal part of Acey_s0009 region corresponding to JARK01001345.1[1,367,393..1,394,816] region, immediately adjacent to AF2 locus. C. LKIN-motif family protein Y032_0009g594 predicted to lay internally in the AF2 locus, between Y032_0009g591 (red oblique font) identified at Acey_s0009 scaffold, corresponding to positions in JARK01001345.1 [1,390,386..1,415,404]. and immediately adjacent, D. LKIN-motif family protein Y032_0009g606 (red oblique font) located in the same panel as LKIN-motif family proteins Y032_0009g610 and Y032_0009g612 located at Acey_s0009 positions corresponding to JARK01001345.1[1,420,759..1,448,188]. E. LKIN-motif family proteins Y032_0009g678, Y032_0009g683 and Y032_0009g686 predicted to lay in the interval between locus AF2 and AB2 on Acey_s0009 at positions corresponding to JARK01001345.1[1,978,782..2,033,870]. F. LKIN-motif family protein Y032_0009g705 predicted to lay internally in the AB2 locus, between Y032_0009g704 (red oblique font) located at the Acey_s0009 corresponding to positions in JARK01001345.1 [2,046,816..2,082,833] and immediately adjacent, G. LKIN-motif protein Y032_0009g716 (red oblique font) located at the scaffold Acey_s0009 at positions corresponding to JARK01001345.1[2,076,229..2,112,237]. The cluster position and the scale (in kb) corresponding to the grid is given in the heading above each panel. Note that genes encoding for the LKIN-motif family proteins do differ in overall length, number and the size of the introns spanning the coding sequences i.e. LKIN-motif protein genes presented at panels F. and G. (both panels are drawn at the same scale indicated by the vertical grid) are different in the overall length by approximately the order of magnitude, while encoded for the relatively small protein products of comparable length.

Based on the above analysis, we note further that the assignment of the LKIN-motif family to the expanded locus on genomic scaffold Acey_s0009 is not absolute. Specifically we emphasize that two other adult specific SL1-capped cDNA clones pAA3 and pAA6 map to scaffolds other than Acey_s0009 (embedding in the expanded AF2-AB2 locus). Instead SL1-capped cDNA clones pAA3 and pAA6 map to at least two different *A. ceylanicum* scaffolds Acey_s0018 (JARK01001354.1) and Acey_s0001 (JARK01001337.1) respectively (Table 2.), suggesting even greater genomic diversity of loci encoding for the LKIN-motif protein family. This perspective is also indirectly supported by our analysis suggesting that the additional locally duplicated clusters of presently uncertain significance, are encoded by scaffolds dispersed across the *A. ceylanicum* chromosomes. Finally, we note, that data summarized in Table 2 indicate that at least two other additional mutually homologous adult specific SL1-capped cDNAs pAE2 and pAG6 map onto the same genomic scaffold Acey_s0009 which includes the expanded locus AF2-AB2 encoding for the LKIN-motif protein family, both supporting the hypothetical protein prediction Y032_0009g563. While presently the function of neither of these factors remains clear, pAG6 appears distantly related to *C. elegans* protein *cri-2,* which is described as a <u>C</u>onserved <u>R</u>egulator of <u>I</u>nnate immunity (Alper et al. 2008). However, this assignment is based on the results of the large scale RNAi screen and therefore requires further verification. Moreover, the significance of the grouping implied by the genetic linkage with LKIN-motif family encoding locus remains presently unclear.

In the broader perspective of the genomic distribution of the *A. ceylanicum* sequences identified by the output from the preliminary characterization, we note that the sequences we generated (table 2) fall into two main categories. The first category involves factors that we mapped with a high stringency search onto genomic scaffolds with a resulting high degree of confidence (described below). The second category involves factors which we could not map unambiguously to specific scaffolds. The second category will not be described in detail herein however the original data are included in Table 2. This second category is not uniform however, since it groups sequences where no genomic scaffold could be assigned (i.e. some of the LKIN-motif members described above), sequences where the relative homology score is weak or coverage is partial and sequences with ambiguous assignments (i.e. where several high scoring matches specify multiple scaffolds). We note that the last category may include some false negatives i.e. as mentioned above, small heat shock protein genes encoded by multiple dispersed paralogous genes, possibly explaining some of the mapping difficulties experienced in the analysis.

Within the set of SL1-capped cDNAs derived from adults, 20 sequences mapped with high confidence to genomic scaffolds (this was supported by typically high homology based search scores, sequence coverage and identity over the discrete region of single specific scaffold) including 5 described above which mapped to the scaffold encoding for the LKIN-motif protein family. Using the same criteria, we have established that 8 SL1-capped cDNAs, derived from the infective larvae, map with high confidence to specific genomic scaffolds other than Acey_s0009. In the case of the set of reads we generated using plasmid clones from genomic DNA, we established that 13 map with a high degree of confidence to specific genomic scaffolds other than Acey_s0009, including a read from the BAC library (supported by the scaffold Acey_s0211). Within that set, two paired reads from genomic DNA clone pVB11, unexpectedly aligned with high confidence to a region of genomic scaffold Acey_s0161 (JARK01001497.1) over the 60-69% of the insert length (Table 2.). The reason why we find this behavior unexpected, is that this clone was previously identified as contamination of mitochondrial origin,and therefore derived from non-chromosomal DNA molecules. Upon close examination, provided the Acey_s0161 scaffold is 476,687kb long, we exclude the possibility that this could represent a mis-annotated mtDNA genome fragment. Detailed alignment identified this 507bp mitochondrial DNA fragment of the insert aligning with high stringency (99% of identity with the DNA alignment) over positions 21,264-21,771 of the genomic scaffold Acey_s0161 (Figure 4.A. and C.). (This was independently confirmed by repeated searches with the entire mtDNA genome of *N. americanus* (Figure 4.B.), and expectedly turned out to be highly conserved with the mtDNA genome of *A. ceylanicum.* Only the above 507bp segment of the mitochondrial genome appears to be included in the specified position of the above genomic scaffold). We interpret this observation as being attributable either to fragmentation of the mitochondrial genome and translocation into the nuclear locus in scaffold Acey_s0161 or alternatively representating an artifactual error of the genomic assembly. If the later possibility can be confidently excluded, the former explanation has precedent in that so called NUMTs (“nuclear mitochondrial DNA segment”)(Lopez et al. 1994) have been described in eukaryotic genomes (Richly and Leister 2004).

**Figure 4.**
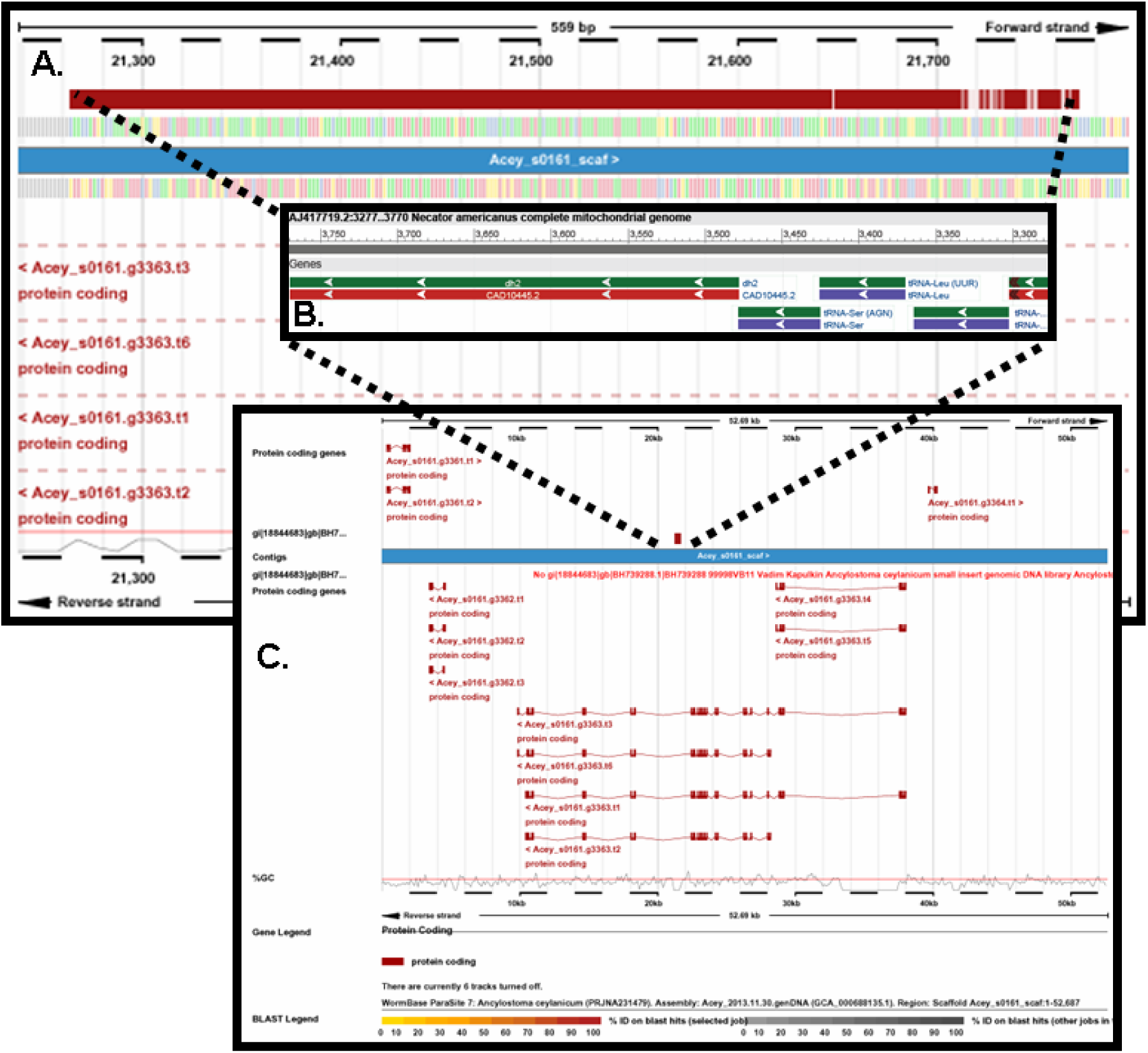
Ensembl representation of pVB11 alignment (the possible nuclear insertion of the mitochondrial segment) onto genomic scaffold Acey_s0161 at position corresponding to JARK01001497.1 [21,264..21,771]. A. *A. ceylanicum* mitochondrial segment 507bp (Ensembl BLASTN match shown as a horizontal red bar, according to color coded BLAST scale) is found inserted into non-alternative intron of the predicted gene Acey_s0161.g3363. B. Corresponding segment of *N. americanus* mitochondrial DNA (AJ417719.2) aligning with same region of the genomic scaffold Acey_s0161 is represented as GeneBank graphics (specific positions of the alignment are given in the image headings). Note fragments of two mitochondrial protein genes at the insert termini and three mitochondrial tRNA genes, present in the segment (position and sequence of all three *A.ceylanicum* tRNA genes identified in the genomic scaffold Acey_s0161 is given in the legend to supplementary Fig.4). C. Position of the intronic insertion of the mtDNA segment is indicated by dotted lines, connecting insertion termini as presented with panel A. and B. (C. is a version of the Ensembl image view of the BLAST alignment shown in panel A., sufficiently enlarged to cover the entire sequence of the Acey_s0161.g3363 gene ~ca 50kb).

**Figure 5.**
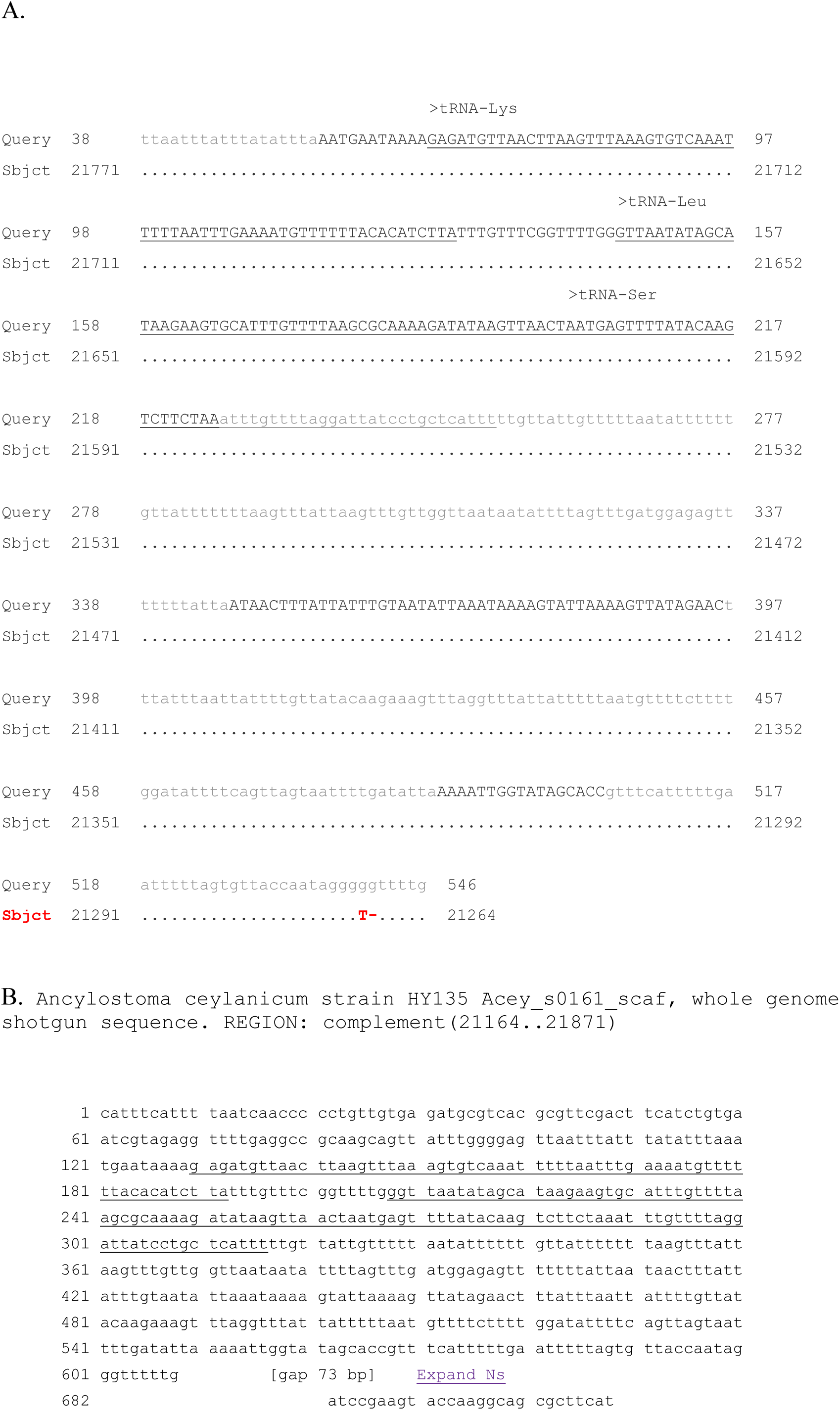
Supplementary information and supplement legend. A. Text view of the pairwise dotted alignment of the query region (729bp) of pVB11 plasmid sequence [gi|18844682|gb|BH739287.1|BH739287 32248 pVB11 Ancylostoma ceylanicum small insert genomic DNA library] against Ancylostoma ceylanicum strain HY135 Acey_s0161_scaf, whole genome shotgun sequence [JARK01001497.1, Length: 476687] resulting in single match on (-) strand in range 1: 21264 to 21771bp, with match alignment statistics (E value 0.0, 507/509 identities). Position and type of the three identified *A. ceylanicum* mitochondrial tRNAs is underlined and tRNA start indicated by ‘>’. B. GeneBank extracted 708bp reverse-complement sequence (embedding the above segment of genomic Acey_s0161_scaffold, with 100bp flanks on each side (25-mar-2014)) with underlined bases corresponding to three mitochondrial tRNA genes. Note the gap of 73bp present at one side of the insert-scaffold junction.

## Discussion

Given the value of recombinant DNA resources, our primary motivation in this study was to develop useful molecular reagents to facilitate subsequent research exploiting transgenic *C. elegans* to study the genes of parasitic nematodes, and notably hookworms. The genetic model organism *C. elegans* is a robust system for the transgenic expression of parasite genes, with numerous genetic tools available, such us RNA interference, fluorescent reporter protein fusions and transgenic complementation (referred as ‘mutant rescue’). At the time when the current project was initiated only a very limited set of hookworms genes had been deposited in the publicly available databases (and these were mostly fragments of mitochondrial DNA and other sequences used for phylogenetic reconstructions).

The resources that we have generated have already contributed greatly to gene discovery in parasitic nematodes (Parkinson et al. 2003), resulting in hundreds of new sequences being deposited in public databases. Since the major resources for gene discovery, the recombinant SL1-cDNA libraries that we generated, have already been described and discussed in some detail (Mitreva et al. 2005), here we will limit the discussion to aspects which have only been discussed poorly so far or have not been described previously at all (recombinants of the genomic DNA).

We note that the approach we implemented for SL1 capped transcripts and for the recovery of plasmid SL1-cDNA clones enables specific identification of the transcripts that engage the nematode specific 5’ end processing referred as trans—splicing (reviewed in, Blumenthal 2012). In nematodes trans-splicing has been mainly implicated in shifts in gene expression associated with the transition of major developmental stages, such as transitions between the successive larval stages or diapausal exit (Zaslaver et al. 2005) and with processing of poly-cistronic transcripts (Blumenthal 2012). The precise physiological role of the 5’ end trans-splicing remains largely elusive and mostly speculative. In *C. elegans* the majority of pre-mRNA transcripts engage in some form of the 5’-end trans-splicing reactions (Blumenthal 2012). While a broad discussion of the role of trans-splicing in developmental transitions and poly-cistronic processing is beyond the scope of this paper, below we concentrate on a few examples directly inferred from our study.

The particular SL1-capped cDNA clone (plasmid pAG1) recovered from the adult stage *Anyclostoma* cDNA library was initially annotated as being homologous with *C. elegans* gene F02A9.2. This SL1 capped cDNA is indeed homologous to each of the locally duplicated pair of *C. elegans* genes F02A9.2/ F02A9.3, listed in Table 2 but not covered in the results section. Mitreva (2005) described a larger set of SL1-capped sequences derived from the cDNA libraries that we had provided (and have described herein), and those transcripts are more correctly classified as encoding for hookworm fatty acid and retinol binding (FAR) proteins. Secreted nematode FAR proteins have drawn the attention of others, due to their postulated role in pathogenesis (Garofalo et al. 2003). In particular, Fairfax et al. (2009) have published on *A. ceylanicum* FAR-1, yet another transcript reported from the larger cDNA set described by Mitreva et al. (2005). We note here that the cDNA that was the subject of those studies, clearly constitutes a different FAR protein from the one described here. The cDNA described in the current work encodes the SL1-capped hookworm FAR protein which is clearly distinct and only partially homologous (approximately 60% identity at the amino acid level) with the one reported by Fairfax et al.(2009). Moreover our SL1-FAR appears to be encoded by different and separate locus (because of their assignment into a different genomic scaffold) in the assembly reported by Schwartz et al. (2015), than the protein described as FAR-1. Nematode FAR proteins are structurally and functionally divergent and therefore typically grouped into clans or subtypes (Garofalo et al. 2003). Therefore it is not surprising in our opinion that detailed analysis has revealed a greater complexity than that previously considered. In the larger SL1-cDNA set (Mitreva et al. 2005) at least twelve other SL1-capped transcripts appear to align with the prototype SL1-FAR sequence that we deposited in the GeneBank database in 2000-2001. Provided that some other cDNAs encoding for FAR proteins exist in the larger set (recovered from the uncapped cDNA libraries provided by other collaborators) and align with the sequence reported by Fairfax et al.(2009) we propose here that the observed SL1 bias represents specific subtypes of the hookworm FAR proteins that will form the basis for future classification of these proteins and their genes.

The set of sequences generated from our recombinant DNA libraries that we have described, includes also another example of an unusual use of the 5’-end transcript processing. This example concerns the SL1-cDNA encoding the hookworm small heat shock protein (plasmid pAF1 in the adult SL1-cDNA set). Small heat shock proteins are encoded by stress sensitive genes and have been extensively characterized in *C. elegans* (Russnak et al. 1983, Stringham et al. 1992, Candido 2002). They form an adaptation to heat stress but also constitute a safeguard against misfolded protein stress (Fonte et al. 2002). Our results suggest that the parasitic adult stages of *A. ceylanicum,* express small heat shock proteins constitutively. This differs from the reported expression pattern of small heat shock protein in *C. elegans* and might reflect an adaptation of the parasitic stages of hookworms to the elevated temperatures that the parasitic stages experience within endothermic hosts. The presence of a trans-spliced SL1 sequence on 5’-end of *A.ceylanicum* messenger RNA encoding a small heat shock protein indicates a radical difference to currently accepted view of the small HSP mRNA in *C. elegans* (where small heat shock protein transcripts are regarded as being excluded from 5’ end trans-splicing). This observation appears to be consistent with what has been reported for other parasitic nematodes i.e. *Nippostrongylus braziliensis* (Tweedie et al. 1993), where SL1 is trans-spliced onto the 5’-end of developmentally regulated small heat shock protein mRNA.

We also note that the anecdotal, proposed association that elevated expression of small heat shock proteins correlates with life-span extension in *C. elegans* (Mendenhall et al. 2012), might also be plausible here (since hookworms are long lived nematodes when compared to *C. elegans* (Beaver, 1988; Plmer 1955; Gems 2000)). In an evolutionary context, the adaptation to elevated temperatures and its possible association of 5’ end trans-splicing with a small heat shock protein gene expression in parasitic nematodes remains a phenomenon of fundamental interest. Adaptation to elevated temperatures has been reported also in tropical isolates of some free-living nematode species i.e. *C. briggsae* but also recently in *Pristionchus pacificus* (Leaver et al. 2016), however none of the free-living species survives at the range of temperatures maintained by the endothermic hosts of adult hookworms. While studies on free-living nematodes were designed to identify the genetic component involved in higher temperature adaptation, no single specific locus has been reported to-date. This suggests that adaptation to elevated temperatures may require highly pleiotropic effects, likely resulting from co-ordinated changes to gene expression at a multiple of levels and involving multiple genes. For example, the genome of *C. elegans* is expected to encode around two-hundred molecular chaperones (Guisbert et al. 2013), including a number of heat shock proteins (Kapulkin et al. 2005). We postulate that the adaptation of certain isolates to elevated temperatures is based on a requirement for global alterations to gene expression and we suggest that alterations to the trans-splicing patterns might fulfill the above adaptive requirement. We propose that some of the above aspects could possibly be studied in transgenic *Caenorhabditis* model. In particular, transgenic lines expressing the hookworm small heat shock protein genes could be assayed for an association of trans-splicing with physiological phenomena related to elevated temperature adaptation, i.e. thermotolerance, malfolded protein response, stress resistance and longevity.

The 5’-SL1 trans-spliced set of sequences described herein includes also SL1-cDNA in which homology to genes known in *C. elegans* could not be detected. Again, the scope of this communication as well as limitations of the manuscript size, precludes us from providing a detailed description of all the factors that were detected. Among the clones sampled there are examples of SL1-cDNAs where some detectable homology is restricted to hookworm species (Table 2.). These genes, presumably hookworm specific, are of great interest, as they may encode novel factors involved in pathogenesis, interactions with the host and host immune response, but also, as taxonomically restricted factors, they may yield the promise of specific therapy targets and perhaps may enable improvement in current therapeutics and control strategies.

From the perspective of hookworm specific genes, collectively we identified thirteen examples of *A. ceylanicum* transcripts within the 5’ trans-spliced cDNA set, where the sequence homology is strictly species specific or where the only homologues are identified in hookworms other than *A. ceylanicum.* Those several examples of the SL1-capped, trans-spliced transcripts (encoding mostly for uncharacterized hypothetical proteins, see results section) could be regarded as examples of SL1– capped transcripts encoding for hookworm specific factors (or possibly events of *A. ceylanicum* specific factors). We emphasize that the retrospective analysis described herein indicates that a number of those hookworm specific factors were detected exclusively within the clones sampled from the SL1-capped cDNA libraries that we provided. Those transcripts encoding for hookworm specific factors were absent from the uncapped cDNA libraries provided by other collaborators. The above perspective, provides an unbiased validation of ours results and clearly and specifically indicates the distinct advantage of the approach we implemented.

Remarkably, when the project was initiated, amongst the hookworm specific SL1-cDNAs at least three identified sequences were found to represent a distinct clade of mutually homologous factors. Among these, homology with other species of nematodes was weak or marginal, confirming that these represent the species expanded gene clade. The homology of the three initially identified SL1-cDNAs (mutually homologous, represented by plasmids pAH6, pAB2 and pAF2) was then extended by the iterated species restricted search. This re-iterated search assigned further pAD5 and pAF6, and pAH1 (in iterated searches on pAH6 and pAB2 respectively), to the hookworm specific SL1-cDNA clade described here, along with three more distantly related SL1-cDNA clones and the number of the re-sampled SL1-cDNAs including the larger set of sequences (Mitreva et al. 2005) generated on SL1-captured clones from the libraries that we had provided. We note that such an extensive representation of the taxon-specific hookworm transcripts (accounting for an estimated ~10-20% of all transcripts) recovered from the relatively small initial set, is possibly indicative of vital relevant physiological function in *A. ceylanicum.* Based on the distinct Leu-Lys-Ile-Asn motif detected in the conceptually translated sequence of the first SL1-cDNA plasmid pAB2, we refer the above polypeptides derived from an adult specific SL1-capped transcripts to the LKIN-motif protein family.

The above clade of the SL1-capped, trans-spliced cDNAs encoding for the LKIN-motif protein family, appears remarkable in the scope and context of the recent assembly of the whole genome assembly of *A. ceylanicum* (Schwartz et al. 2015). While loci encoding for SL1-cDNAs, pAB2 and pAF2 (and further pAH6) are clearly identifiable within the set of genomic scaffolds, in contrast genes encoding for the SL1-cDNA pAH1 (and further pAD5 and pAF6) do not seem to exist within the preset whole genome assembly of the *A. ceylanicum* genome. This strongly suggests that the present whole genome assembly is not entirely complete (although we note here that the present assembly provided us with fair representation across all other factors listed in Table 2.). Curiously, mapping of the above *A. ceylanicum* trans-spliced SL1-capped transcripts, has demonstrated that, where genomic scaffolds could be attributed, the above factors appear to be linked on the single genomic scaffold Acey_s0009 (with the locally duplicated gene pairs spaced at ~1MB located in the middle of (3,042 546 MB) scaffold). While a detailed description of the complexity at the locus encoding the species-specific trans-spliced hookworm factors was covered only superficially in the result section (more comprehensive description will be submitted under separate cover), we note here that based on our results, the above clade must represent a family of proteins encoded by genes subjected to local chromosomal expansion. In view of the considerations given above about genes subjected to local chromosomal expansion, we foresee that these genes, encoding for related, adult hookworm specific factors, encoded in a relatively narrow genetic interval, might be an example of a metazoan form of a pathogenicity island, as described in pathogenic prokaryotes (i.e. reviewed Schmidt & Hensel 2004). Given the obligatory parasitic lifestyle of adult stages of hookworms, we speculate that these abundant, presumably secreted, factors are involved in aspects of the host-parasite relationship that are relevant in pathogenesis. Hookworms are known to secrete copious amounts of immunogenic materials, including small proteins, and we suggest that the highly diverged members of the LKIN-motif protein family encoded by multiple expanded loci in the hookworm genome (including that described above on scaffold Acey_s0009 and at least two other loci referred in the results section) might collectively act as anti-immunological smokescreen (See Kemp et al. 1987, Ahlborg et al. 2002)diverting the host immune response away from crucial antigens that if attacked by the immune system would disable the parasite. We further propose that highly diverged members of the LKIN-motif family engage the attention of the host immune system by exposing the secreted cloud of partially overlapping, related (but distinct and not identical) epitopes to create immunological mischief leading to the successful immune evasion and hence chronic survival of the parasite..

Another issue regarding the above clade of trans-spliced hookworm specific proteins which became apparent during sequence analysis, concerns a similar sequence reported by Bungiro et al. (2002). Those authors described one of the sequences originating from our SL1-cDNA libraries, as the ‘Excretory/Secretory protein 2’. Bungiro et al. (2002) indicated that the protein corresponding to single EST deposited in GeneBank is found in the material shed by *A. ceylanicum* adults and that the relevant gene product engages the humoral antibody response in laboratory hosts. However, these authors failed to specify the source and origins of the SL1-cDNA sequence (instead, these authors, acknowledged the names of 35 other collaborators). In conclusion, we note that the SL1-cDNA plasmids that we generated and provided through collaborators to the scientific community could possibly have the following practical application. Provided that the SL1 capped full length cDNA are selected by the procedure that we implemented, and given that the vector sequence provides T7 polymerase binding sites immediately adjacent to SL1-cDNA cloning site, we suggest that the plasmids from the libraries we provided are suited as templates for *in vitro* transcription of the messenger RNAs (i.e. constitute material directly suitable for use as the so called mRNA vaccines i.e. Weiner 2013). Otherwise these and other recombinant DNA libraries we have constructed may serve as useful reagents for the T7 polymerase dependent *in vivo* synthesis of double-stranded RNA synthesis in *E. coli* (Timmons et al. 1998), established as a mediator in RNA silencing and RNA interference (Fire et al. 1998). Libraries of the plasmid clones expressing dsRNA are established screening reagents applied widely in *C. elegans* (i.e. relevant references in Kapulkin et al. 2005) and it is therefore expected that these tools could be applied with *C. elegans* lines expressing transgenic hookworm genes.

So far we have generated and described two types of recombinants (*E. coli* replicons) with inserts derived from genomic *A. ceylanicum* DNA: a small insert library and a Bacterial Artificial Chromosome library. Genomic DNA libraries comprise a valuable resource for follow-up analysis in transgenic *C. elegans.* First, gene expression controlling regions could be recovered and assayed with reporter genes (Chalfie et al. 1994). Screens, such us ‘promoter-trapping’ were established to identify *de novo* regions controlling gene activity. Alternatively, candidate genes could be replaced or tagged in genomic constructs to form functional chimaeric reporter genes, suitable for testing in transgenic *C. elegans.* Second, gene encoding sequences in genomic DNA, which are interrupted by introns, could be assayed in transgenic *C. elegans* for correct splicing in cis-but also trans-splicing assays. Third, functional genes of interests could be introduced into *C. elegans* as transgenes, and assayed for expression and specific functions. Together, the constructed libraries of genomic DNA of *A. ceylanicum,* should suit all the above purposes.

During the process of construction of genomic DNA recombinants a few nuances became apparent which we briefly discuss below. The sequence tags generated on the plasmid clones recovered from the small insert library included mitochondrial DNA fragments. While at the time these were interesting due to their conserved synteny with other nematodes, these fragments are presently regarded as unwanted contamination. The particular fragment of the *A. ceylanicum* mitochondrial DNA we described, is unexpectedly found in the nuclear genomic scaffold Acey_s0161 (Schwartz 2015). The inserted segment of the mitochondrial DNA includes fragments of two mitochondrial protein genes and three mitochondrial tRNA sequences, located in the non-alternative intron of *A.ceylanicum* gene Acey_s0161.g3363. We note that it is presently uncertain whether this sequence can be regarded as NUMTs or whether it represents a whole genome assembly artifact. NUMTs have not been previously reported in hookworms. Nevertheless, it is interesting that in filarial nematodes non-chromosomal DNA (i.e. DNA originating from endosymbionts) has been found in the nuclear genome (Dunning Hotopp et al. 2007, McNulty et al. 2010). While hookworms, including *A. ceylanicum* are thought not to have prokaryotic endosymbionts, we think it would be interesting if the NUMTs in *A. ceylanicum* were to have been acquired via a similar mechanism to that responsible for segments of *Wolbachia* DNA being acquired into filarial chromosomes. We note however that the incontroversial demonstration of the potential existence of NUMTs in the hookworm genome will require further investigation. We note also that NUMTs pose a specific challenge with *de novo* whole genome assembly, as mitochondrial DNA could easily contaminate raw nuclear DNA preparations used for PCR based whole genome sequencing. Indeed mtDNA contaminations could be avoided if more careful size exclusion DNA fractionation were to be exploited (Osoegawa et al. 2001).

The other aspect that we feel needs addressing is cloning efficiency. Based on a positive experience with the RT-PCR amplified SL1-cDNA, we were encouraged with the commercially provided vaccinia virus topoisomerase system (see materials and methods), which consistently yielded a low level of non-recombinant clones. However when we attempted to use a parallel product from the same supplier, we were discouraged with an unexpectedly high background of non-recombinant colonies or mis-recombinant plasmids (detected with sequencing, not shown). We initially attributed this unwanted and unforeseen obstacle to some intrinsic properties of hookworm DNA. However our later experience with constructing other plasmid libraries entailing inserts derived from hookworm DNA, with conventional T4 ligase and restriction enzymes, suggested that in the above instance, the supplier provided a vaccinia virus topoisomerase based on the TOPO-plasmid ligation system that performed sub-optimally. While, detailed consideration of the reasons for this dysfunctional performance are outside the scope of this paper, we consider that the overall kinetics of the blunt end (versus T/A) topoisomerase mediated insert-plasmid ligation was impaired or affected by other factors significantly reducing its efficiency.

Despite the above limitations, we have nevertheless managed to construct a Bacterial Artificial Chromosome library in a low-copy vector. Bacterial Artificial Chromosmes (BAC) are large plasmids derived from the bacteriophage P1 or *E. coli* F factor plasmid (Osoegawa et al. 2001) and have proved to be supreme tools in modern genomic analysis. BACs incorporate recombinant DNA fragments sufficiently large to detect haplotype differences over fingerprinted maps of the genomic regions. The so called paired-end sequences of BAC clones are important in genomic reconstruction of chromosomal regions of interest, as the paired-ends provide higher order contig/scaffold coordinates. We have described the construction of a fosmid size recombinant DNA library of *A. ceylanicum* using the BACPAC-CHORI protocol, with slight modifications as applied in another parasitic nematode (Jeremy Foster, NEB, personal communication (Foster et al. 2004)). Importantly BAC accommodated genomic DNA inserts, could be modified via DNA engineering using recombination in *E. coli* methods (Zhang et al. 1998) rendering the recombined clones suitable for expression analysis in transgenic *C. elegans* (i.e. Sarov et al. 2006). While we are aware that the constructed *A. ceylanicum* BAC library referred to above, is limited by insufficient characterization at the sequence level, we nevertheless consider that the available permanent recombinant DNA resource will provide a valuable reference for an improved genome assembly as well as for the recombinant platform for future gene expression studies in transgenic *C. elegans.*

## Author’s contribution

WJK and HW conceived, designed and implemented the experiments and analyzed the data, AM assisted with SL1-library preparation and arraying the plasmid clones, AC performed PFGE DNA fractionation, ML perfomed sequencing on the BAC clone. WJK and JB drafted a manuscript. HW (Parasitology Department Chair) managed the awards and grants. WJK covered the costs of the external sequencing service (www.microsynth.ch).

## Acknowlegements

We thank Ludger Altrogge and Ruth Chiquet (Friedrich Miescher Institute, Basel Switzerland) for introducing us to vaccinia virus based topoisomerase plasmid construction system (Invitrogen) and supporting Wadim Kapulkin visit to FMI in 1998 (respectively). We thank Pieter De Jong (BACPAC resources CHORI, Oakland) for providing pBACe3.6 vector and advice on Bacterial Artificial Chromosome library construction. We are indebted to Christopher Link and Patricia Tedesco (Institute for Behavioral Genetics, Colorado University-Boulder) for suggestions with an initial version of the manuscript (2000-2002). The *Ancylostoma ceylanicum* strain used in this work (referred as ‘Warsaw strain’) was originally provided by Jerzy Behnke (Nottingham University) and was maintained by Ewa Janecka at the Faculty of Veterinary Medicine in Warsaw. The animal studies were approved by the Animal Care and Use Committee (R-508/98). The work was partly supported by the KBN grant nr 5PO6K 006 15 and the special award BNW-N-Vb-56/01 of Faculty of Veterinary Medicine, Warsaw to Wadim Kapulkin (to cover the costs of the outsourcing of the sequencing reactions with external facility service).

